# Ngn2 induces diverse neuronal lineages from human pluripotency

**DOI:** 10.1101/2020.11.19.389445

**Authors:** Hsiu-Chuan Lin, Zhisong He, Sebastian Ebert, Maria Schörnig, Malgorzata Santel, Anne Weigert, Wulf Hevers, Nael Nadif Kasri, Elena Taverna, J. Gray Camp, Barbara Treutlein

## Abstract

Human neurons engineered from induced pluripotent stem cells (iPSCs) through Neurogenin 2 (Ngn2) overexpression are widely used to study neuronal differentiation mechanisms and to model neurological diseases. However, the differentiation paths and heterogeneity of emerged neurons have not been fully explored. Here we used single-cell transcriptomics to dissect the cell states that emerge during Ngn2 overexpression across a time course from pluripotency to neuron functional maturation. We find a substantial molecular heterogeneity in the neuron types generated, with at least two populations that express genes associated with neurons of the peripheral nervous system. Neuron heterogeneity is observed across multiple iPSC clones and lines from different individuals. We find that neuron fate acquisition is sensitive to Ngn2 expression level and the duration of Ngn2 forced expression. Our data reveals that Ngn2 dosage can regulate neuron fate acquisition, and that Ngn2-iN heterogeneity can confound results that are sensitive to neuron type.

## Introduction

Human cell types engineered from induced pluripotent stem cells (iPSCs) through transcription factor overexpression are widely used to study the mechanisms controlling cell fate differentiation, to model human diseases, and to identify potential therapies (Guo and Morris, 2017). Human neurons can be generated through the forced expression of the transcription factor Neurogenin 2 (Ngn2) with high efficiency and reproducibility (Zhang et al., 2013). These Ngn2-induced neurons (Ngn2-iNs) functionally mature into morphologically complex and electrophysiological active neurons after approximately 3-4 weeks of co-culture with astrocytes. The Ngn2-iN system has been used extensively to understand neuron development and model disease (Aneichyk et al., 2018; Lin et al., 2018; Meijer et al., 2019; Pak et al., 2015; Yi et al., 2016). However, the characterization of Ngn2-iNs so far has generally been limited to functional assays, biomarker expression, and bulk transcriptomics. There is lack of comprehensive transcriptomic comparison with primary neuron subtypes and it is unclear whether any off-target fate emerge during the differentiation process. Single-cell sequencing methods provide powerful resolution into the heterogeneity of directed differentiation culture systems (Biddy et al., 2018; Cacchiarelli et al., 2018; Camp et al., 2018; Karow et al., 2018). Previously, we have used single-cell mRNA-sequencing (scRNA-seq) to dissect the differentiation path from mouse embryonic fibroblasts and human percityes to neurons and identified previously undescribed heterogeneity generated by the overexpression of the pioneer factor Ascl1 (Karow et al., 2018; Treutlein et al., 2016). Here we set out to characterize Ngn2-iNeuron heterogeneity, identify the cell states that are generated during differentiation, and analyze the dynamics of the differentiation process using scRNA-seq.

## Results

### Heterogeneity of Ngn2-induced neurons dissected by scRNA-seq

We generated a stable iPSC line with lentiviral insertion of two constructs, one constitutively active rTet under a constitutive EF1a promoter and the second expressing Ngn2 under a tetO promoter. In this way, Ngn2 can be induced by Doxycycline (Dox), which drives the differentiation towards iNeurons (Frega et al., 2017; Zhang et al., 2013). We then performed high-throughput droplet microfluidic scRNA-seq (10X Genomics) that is based on 3’ transcript counting with unique molecular identifiers (UMIs) at multiple time points during directed differentiation (Fig. 1A). After filtering the data of astrocytes, multiplets, and cells with insufficient UMIs, a total of 6,764 cells (d0, 1,412 cells; d1, 2,688 cells; d5, 524 cells; d14, 1,515 cells; d28, 625 cells) were included in the analysis.

**Fig. 1.**
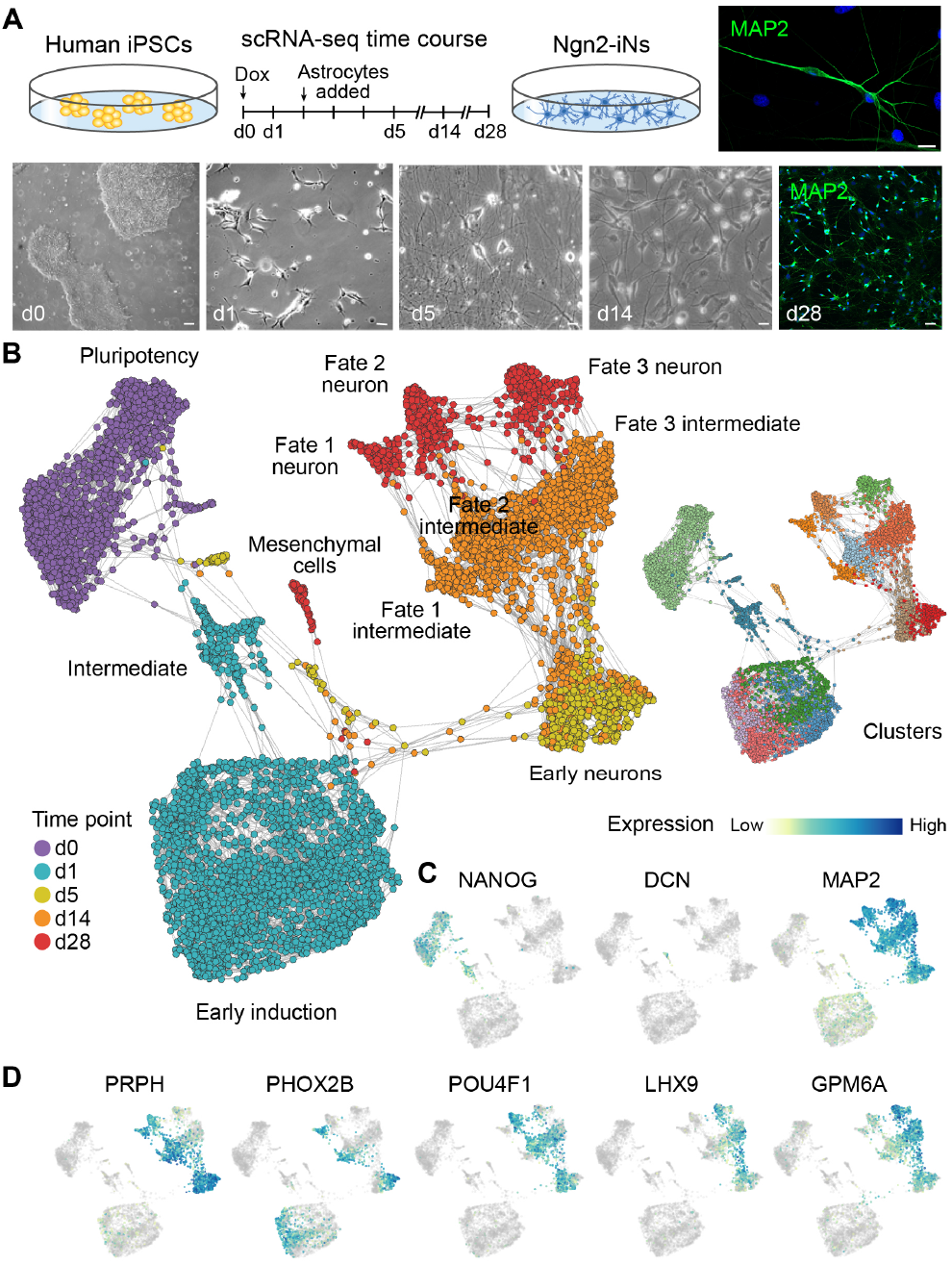
Diverse subpopulations emerge during Ngn2-directed neuron differentiation into iNs from human iPSCs. (A) Schematic of scRNA-seq time course experiment and representative images from human iPSCs differentiating into Ngn2-iNs. Polyclonal human iPSCs (409b2) stably integrated with Doxycycline-inducible Ngn2 were directly differentiated into neurons. Cells were analyzed with scRNA-seq (10X Genomics) at multiple time points during differentiation (d0, d1, d5, d14, d28). Rat cortical astrocytes were added to promote neuronal maturation and synaptic connectivity. Immunohistochemical staining of Ngn2-iNs at d28 with MAP2 (green) and DAPI (blue). Scale bars are 10 μm. (B) SPRING embedding shows the developmental relationships of 409b2-derived Ngn2-iNs with cells colored by time points (left) or cluster (right). (C-D) Expression feature plots of pluripotency marker (NANOG), mesenchymal marker (DCN), and pan-neuronal marker (MAP2) are shown (C) as well as markers for Ngn2-iNs subtypes (D).

To reconstruct the differentiation path of Ngn2-iNs, we combined all of the time course data, identified clusters (Stuart et al., 2019), and visualized cell relationships in a 2D graph embedding (Weinreb et al., 2018) based on genes that vary at each time point (Fig. 1B, C and Fig. S1A). Surprisingly, we found at least 4 transcriptionally distinct cell populations at the d28 time point. One population is marked by DCN, COL5A1, and other extracellular matrix proteins and we interpret this cluster as an off-target mesenchymal population. In addition, we observed three different neuronal clusters that express high levels of pan-neuronal genes (MAP2, NCAM1) yet are molecularly distinct. Two clusters have high expression of PRPH, a gene that is highly expressed in neurons of the peripheral nervous system (Yuan et al., 2012). These two PRPH+ clusters segregate into a PHOX2B+ cluster and a POU4F1+ cluster (Fig. 1D). The other neuronal cluster is marked by GPM6A expression, which is expressed throughout both the central nervous system (CNS) and the spinal cord during mouse development (Fig. 1D) (Diez-Roux et al., 2011).

Ngn2 induction resulted in major gene expression changes early on in programming (d0, d1 and d5), likely driven by the immediate downstream targets of Ngn2. This transition was accompanied first by a decrease in NANOG, which was downregulated already by d1, followed by subsequent decrease of POU5F1, indicating loss of pluripotency. Meanwhile, MAP2 expression increased starting from d1, suggesting rapid induction and development into the neural lineages (Fig. 1B and Fig. S1B). Due to this rapid transition from iPSC to cells committed to a neuronal fate, we hypothesized that directed differentiation bypasses early transitional states that are usually observed *in vivo* to reach neuronal states. To test the hypothesis, we ordered Ngn2-iNs in pseudotime based on transcriptome similarities and compared the resulting trajectories with development of neurons in brain organoids, which progress step-wise through neuroectoderm, neuroepithelium, and neural precursor stages from pluripotency (Fig. S1C-D) (Kanton et al., 2019). We observed an escape from the early developmental stages from pluripotency directly into neural precursor stages, skipping multiple intermediate stages including neuroectoderm and neuroepithelium induction, supporting a more direct differentiation model (Fig. S1D).

To investigate the molecular events underlying the dramatic developmental changes, we identified 3,231 genes with significant expression changes along the course of Ngn2-iN development and segregated them into six clusters with their expression peak at different stages (Fig. S1E). We performed functional enrichment analysis on these differentially expressed (DE) genes and recovered gene ontology terms related to neural development, highlighting changes from proliferation, morphogenesis, synaptogenesis to functional synapses formation (Fig. S1F and Supplementary Table 1). We cross-referenced DE genes with annotated transcription factors (TFs) (Hu et al., 2019) to identify potential drivers of gene expression changes during Ngn2-iN development (Fig. S1G) (Hu et al., 2019). Based on the observation of early neural induction, we focused on TFs that changed from 6-12h to d1 after Dox induction and constructed a gene regulatory network (Aibar et al., 2017), incorporating transcription factor binding site prediction in promoters with TF-target co-expression (Fig. S1H) (Aibar et al., 2017). Notably, although Ngn2 was not included in the network due to its constant expression in the system, it was predicted to connect with some of TFs with the highest centrality in the constructed regulatory network, including POU5F1, HES6 and SOX11, supporting its role in driving direct reprogramming from iPSCs to induced neurons.

While neural induction begins before d1, the heterogeneity of Ngn2-iN emerges later as the expression of Ngn2-iN subtype markers PHOX2B, POU4F1 and GPM6A were detected after d1 (pseudotime Pt 0.4) (Fig. S1B and I). Interestingly, we found that PHOX2B and POU4F1 had divergent expression from the beginning of their activation, while POU4F1 and GPM6A bifurcated later at d5 (pseudotime Pt 0.75) (Fig. S1J and K). PRPH, on the other hand, was detected only after the PHOX2B and POU4F1 bifurcation, suggesting that it was activated independently in PHOX2B and POU4F1 expressing cells (Fig. S1I).

### Ngn2-iN heterogeneity is commonly detected

We next determined if the heterogeneity of Ngn2-iNs results from heterogeneous iPSC populations used to induce iNs, or if the heterogeneity was specific to the particular iPSC line. We therefore established a single iPSC clone (409b2 monoclonal) from the parent 409b2 line used in the time course data, and we additionally generated a polyclonal Ngn2-inducible line from another individual (Sc102a1). We induced Ngn2 polyclonal 409b2, monoclonal 409b2, and Sc102a1 lines (Fig. 2A) and analyzed the resulting transcriptomes at d35 of differentiation (Fig. 2B). We found that significant heterogeneity is still observed in an integrated analysis of the scRNA-seq data, with 6 molecularly distinct clusters containing cells from each of the starting iPSC lines. We note that the three main neural subtypes GPM6A+, PRPH+/PHOX2B+, and PRPH+/POU4F1+ are observed for each line, as well as off-target cells (Fig. 2B-E). We searched for the top differentially expressed genes in the 6 clusters, and observed distinct gene expression patterns for each cluster (Fig. 2F). These data suggested that Ngn2-based neural reprogramming is intrinsically heterogeneous, independent of the clonality of the starting cell population, and that the heterogeneity is detected in neurons from multiple iPSC lines.

**Fig. 2.**
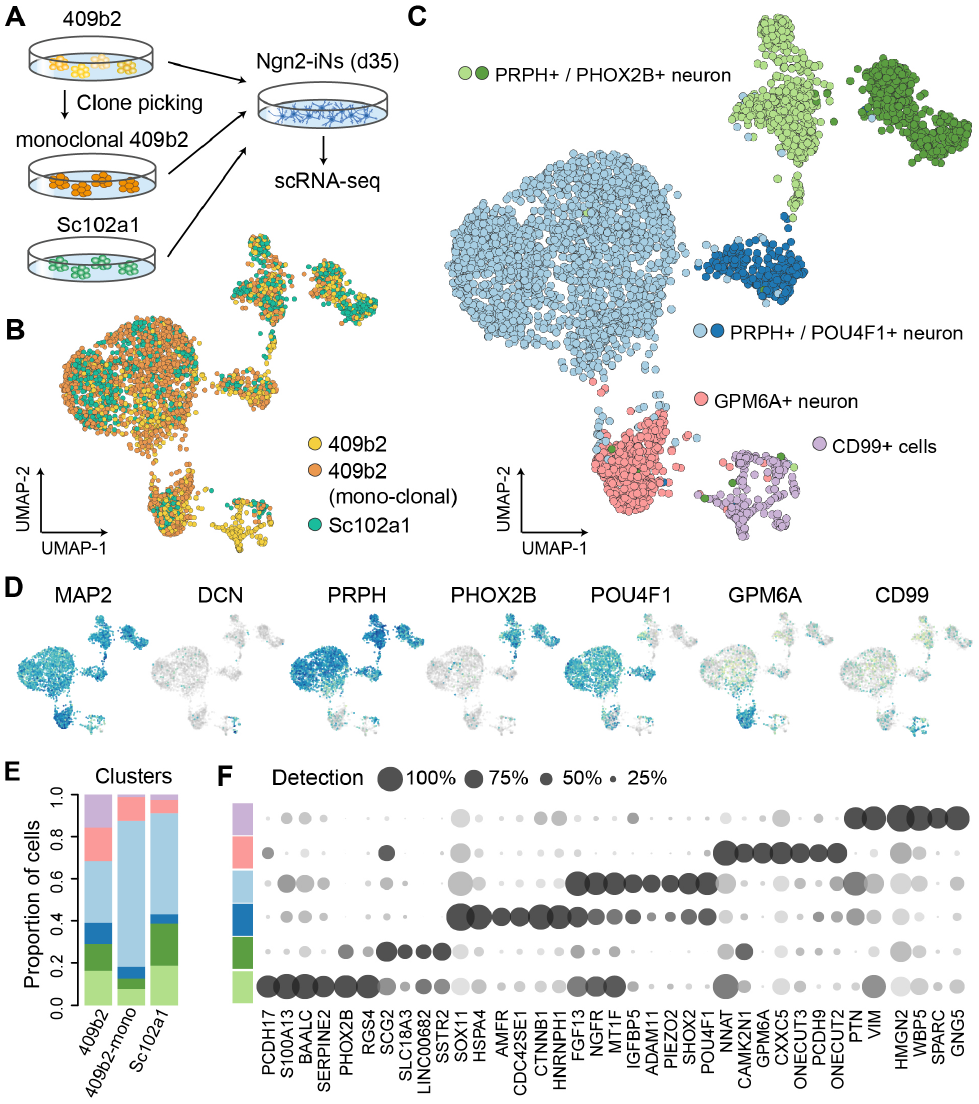
Ngn2-iN neuron diversity is recapitulated in multiple iPSC lines. (A) scRNA-seq was performed on d35 Ngn2-iNs from polyclonal 409b2, monoclonal 409b2 and polyclonal Sc102a1 iPSCs. (B-C) UMAP embedding of Seurat 3.0 integrated scRNA-seq data, with cells colored by cell source (B) or cluster annotated by marker genes (C). (D) Feature plots showing the expression of marker genes. (E) Stacked bar plot showing proportions of clusters in each sample. (F) Dot plot of marker gene expression patterns and detection rates across clusters.

### Molecular features of Ngn2-iN subpopulations

We next analyzed the molecular signatures that distinguished each Ngn2-iN cluster in order to better understand the identity of each population. Towards this aim we compared the signatures to in situ hybridization (ISH) data from the Allen Developing Mouse Brain Atlas, and to single-cell transcriptome atlases from the mouse central and peripheral nervous system and human cortex and retina (Fig. 3A). We also assessed the expression of neurotransmitters and other markers of neuron specialization. Based on whole-embryo mouse ISH data (Thompson et al., 2014), we find that *Ngn2* is expressed in progenitor zones in the developing telencephalon, however it is also detected in many other brain structures including derivatives of the dien-, mesen-, and rhombencephalon, as well as undefined structures in the spinal cord (Fig. 3B and Fig. S2). *Prph* is expressed in the neural retina, trigeminal nerve, and nuclei within the grey horn of the spinal cord. *Phox2b* is expressed in rhombencephalon/brain stem neurons as well as neurons in the PNS. *Pou4f1* is expressed in the retina, mesencephalon derivatives, trigeminal nerve, and grey horn nuclei. We note that *Phox2b* and *Pou4f1* are anticorrelated, and Phox2b has been demonstrated in regulating a switch from somatic to visceral sensory neuron identity (D’Autreaux et al., 2011). We next compared each Ngn2-iN cluster to PNS and CNS neurons from primary reference cell atlases (Clark et al., 2019; La Manno et al., 2020; Zeisel et al., 2018) (Fig. 3C). Unlike neurons in the iPSC-derived cerebral and retinal organoids, the Ngn2-iN clusters did not show specific substantial transcriptomic similarity to any CNS neuron subtypes. Meanwhile, they were relatively similar to PNS neurons, especially the PRPH+ clusters, although they were not specifically similar to any of the PNS neuron subtypes. We next explored if Ngn2-iN expressed markers of the primary neuron subtype (Fig. 3D and Fig. S2). Though there is some overlap, we could not identify a clear cell *in vivo* neuron population that was obviously the counterpart of any Ngn2-iN cluster. Furthermore, based on the PNS and CNS neuron average transcriptomic profiles in the primary references, we deconvoluted the ratio of PNS/CNS identity for each cluster. We found that all Ngn2-iNs have mixed signatures of CNS- and PNS-derived neurons, and do not show a clearly established identity based on this metric, whereas primary human CNS or PNS neurons, as well as human cerebral organoid neurons, have more clearly distinguished biases (Fig. 3E). We find that the GPM6A+ cluster has more CNS features while the PRPH+ clusters have a biased PNS signature, in line with GPM6A and PRPH showing high expression in CNS and PNS neurons, respectively. Altogether, our data suggested that Ngn2-iNs have a mixture of neuronal signatures, and we were not able to establish a clear identity of Ngn2-iN populations. We note that this lack of *in vivo* counterpart could be due to incomplete reference cell atlases, as well as discrepancies between human and mouse neurons.

**Fig. 3.**
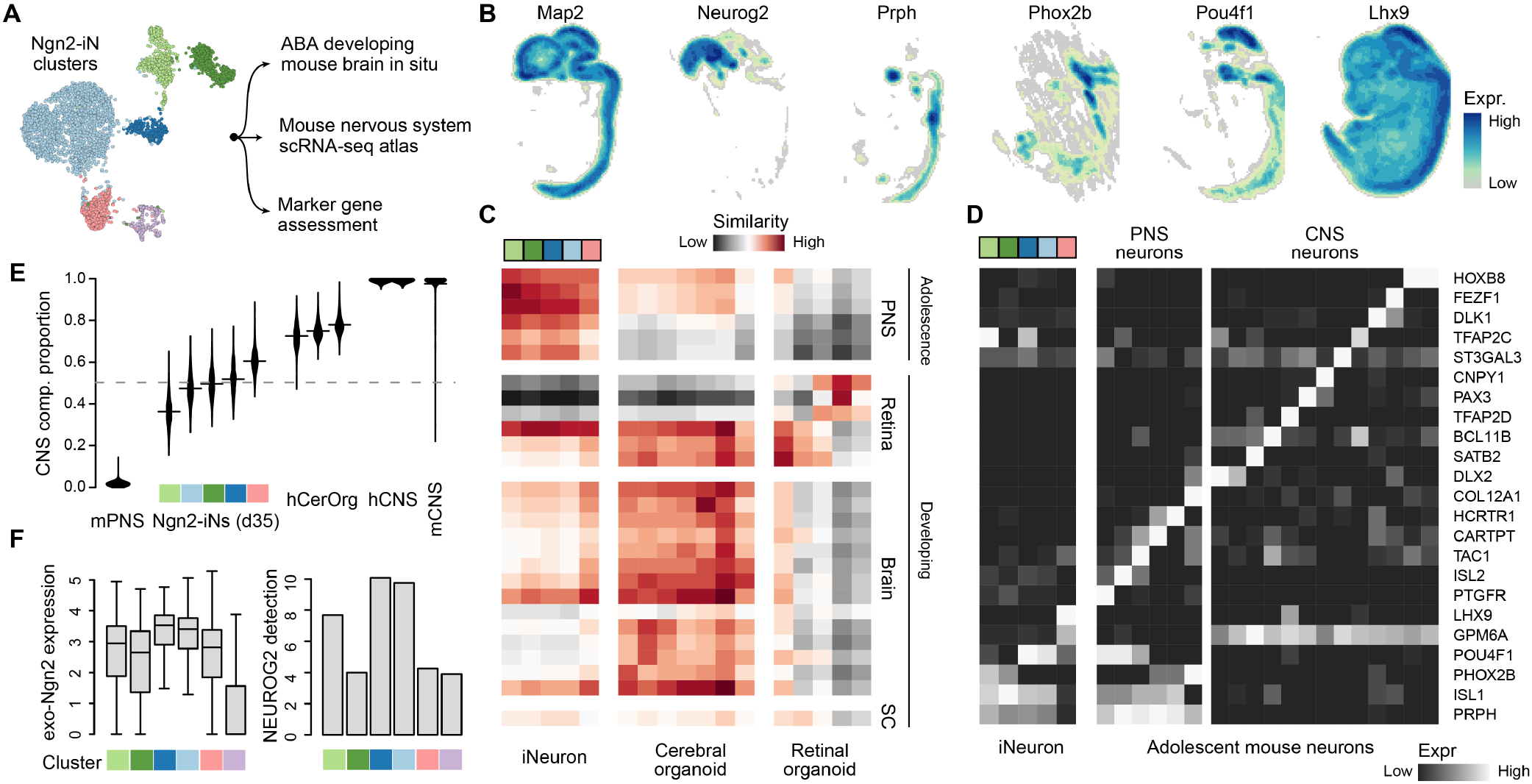
Molecular signatures of Ngn2-iN compared to primary neuronal cell types mouse reference atlases. (A) Ngn2-iN subpopulation signatures were compared to diverse reference atlases. (B) Spatial expression patterns of selected markers as maximum intensity projections across sagittal sections in the E13.5 mouse brain from the Allen Developing Mouse Brain Atlas. (C) Transcriptomic similarities between iPSC-derived neurons (Ngn2-iNs, cerebral organoids (Kanton et al., 2019), and retinal organoids (Cowan et al., 2019)) and primary mouse neuron subtypes (Clark et al., 2019; La Manno et al., 2020; Zeisel et al., 2018), represented as Pearson correlations between expression profiles. (D) Average expression of various marker genes of primary neuron subtypes in Ngn2-iN clusters and primary mouse PNS and CNS neuron subtypes. (E) Proportions of the estimated CNS component in Ngn2-iNs, cerebral organoid neurons, and human/mouse primary mature PNS/CNS neurons. (F) Expression of exogenous (left) and endogenous (right) Ngn2 in different Ngn2-iN clusters. The expression of exogenous Ngn2 is represented by log-normalized UMI, and the expression of endogenous NEUROG2 is represented by detection rates.

The expression level of reprogramming factors could affect the outcome of reprogramming (Sommer et al., 2012) and lead in part to the heterogeneity Ngn2-iNs. We thus analyzed the relationship between Ngn2 expression and the molecular identity of corresponding Ngn2-iNs. The Ngn2 expression level and proportion of Ngn2-expressing cells was indeed lower in the off-target cluster (CD99+), in line with previous studies that failed reprogramming is linked to silenced reprogramming factors (Treutlein et al., 2016) (Fig. 3F). Among the successfully reprogrammed neural clusters, we observed variable expression levels of Ngn2 and proportion of Ngn2-expressing cells, prompting us to examine whether duration of Ngn2 induction affects the Ngn2-iN reprogramming heterogeneity.

### Duration of Ngn2 induction affects Ngn2-iN subtype configuration

We manipulated Ngn2 expression by shortening the duration of Dox treatment, from continuous treatment throughout the course of induced-neural development to removal after day 1, day 3 and day 5 of treatment, and analyzed the resulting cells at day 14 using scRNA-seq (Fig. 4A). A total of 2767 cells (d1, 311 cells; d3, 378 cells; d5, 727 cells; d14, 1351 cells) were included in the analysis. As expected, the expression level of exogenous Ngn2 is correlated with the duration of Dox treatment (Fig. 4B). Endogenous NGN2 expression is also positively correlated with Dox treatment duration, likely as a result of the positive autoregulation (Fig. 4B) (Ejarque et al., 2013; Quinones et al., 2010). Each of the previously identified major clusters were detected in all samples, however the duration of Dox treatment affects the proportion of samples among each cluster (Fig. 4C-E). Specifically, we found that the GPM6A+ population was enriched in samples with shorter Dox treatment, while the PRPH+/POU4F1+ population was more abundant in samples with increased Dox treatment (Fig. 4E). We found that the proportion of PHOX2B+ cells was not affected, which could be due to the observation that PHOX2B expression is activated early in reprogramming (day 1 after Ngn2 induction), and this fate trajectory could be established before we manipulate the Ngn2 expression (Fig. S1J). This data shows that the duration of Ngn2 induction impacts the proportion of neuron subtypes that emerge in this single factor reprogramming paradigm.

**Fig. 4.**
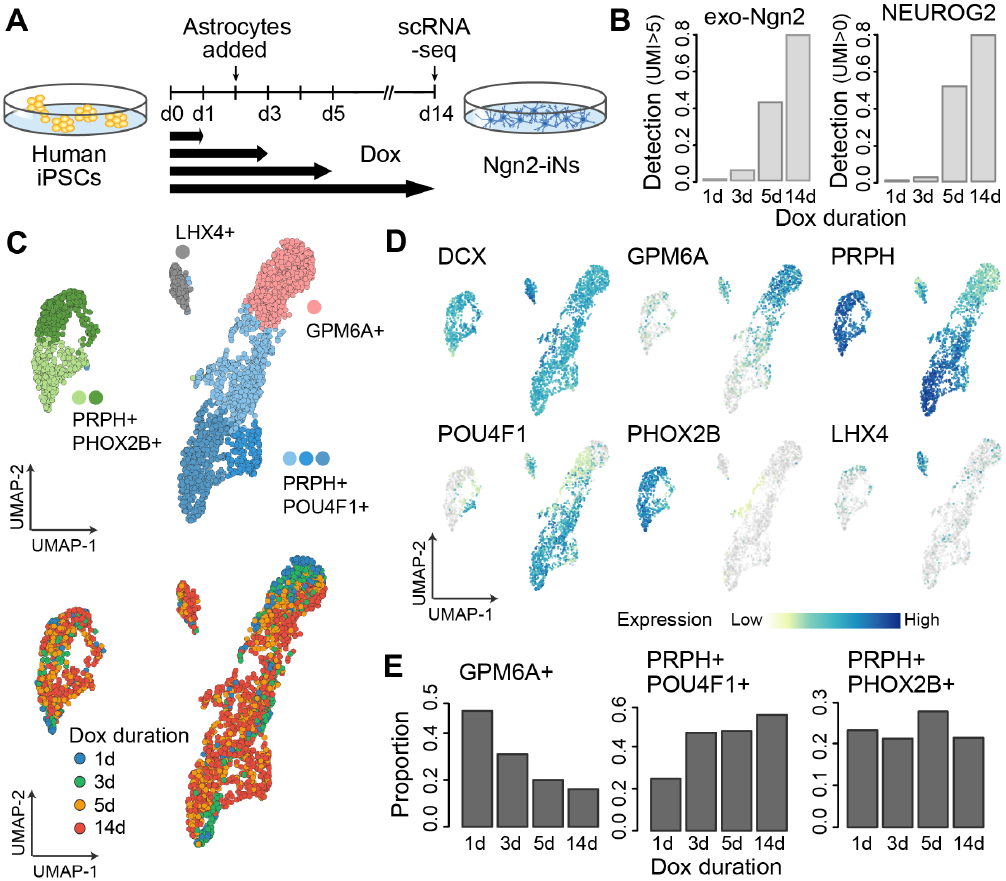
Neuronal fate specification is sensitive to duration of Ngn2 induction. (A) Schematic for the Dox treatment duration experiment. Ngn2 expression was induced with Dox in 409b2 iPSCs at d0. Dox was then removed at d1, d3, d5 and d14. Corresponding Ngn2-iNs were collected for scRNA-seq at d14. (B) Detection rates for cells expressing exogenous Ngn2 (>5 UMI) and endogenous Ngn2 (>0 UMI) from samples in (A). (C) UMAP analysis of Ngn2-iNs from experiment in (A). scRNA-seq data were integrated using cluster similarity spectrum (CSS)-based integration (He et al., 2020). Cells were colored by cell sources (top) and UMAP clusters (bottom). (D) UMAP plots colored by marker gene expression. DCX marks the neural lineages. LHX4 marks the identity of the off-target cluster as the immature cone photoreceptor (Buenaventura et al., 2019). (E) Proportion of cells from samples in (A) in the three neural clusters.

## Discussions

Cell fate engineering of neural subtypes from human iPSCs using defined transcription factors provides extraordinary new inroads into disease modeling and therapy screening using human cells. Methods to rapidly generate mature human neurons are exciting and transformative for these endeavors. There are many scenarios where neurons with generic morphological features or electrophysiological activity are sufficient, and indeed the Ngn2-iN system has been a valuable tool in such cases. However, there are physiological and disease scenarios where it will be critical to engineer precise neuron cell states with high efficiency. Our analysis of the single-factor Ngn2 overexpression system using permissive culture conditions suggests that the emergent neuron population is heterogeneous, with the heterogeneity being consistent across different cell lines. We are unable to assign the neuron populations to a particular identity with high confidence. We note that this may be due to the fact that current single-cell and spatial transcriptome reference atlases are incomplete. However, without specific matrix and guiding molecules it may be expected that neurons are not able to establish molecular profile observed *in vivo* with high precision. We also highlight that Ngn2 is dynamically and variably expressed in the developing mammalian central nervous system, supporting that forced Ngn2 expression could result in diverse neuronal populations. In fact, our data shows that multiple Ngn2-iN subpopulations are more similar to neurons of the peripheral nervous system than they are similar to the central nervous system, and it is unclear if this culture paradigm is indicative of CNS functionality. A further confounding factor in this system is that the lentiviral construct containing the Ngn2 cassette can become silenced, resulting in stochastic aberrant neuronal states. Indeed, our data suggests that Ngn2 expression level and timing of overexpression has a major role in defining t he n euron s tates t hat are generated. It is a challenging endeavor to engineer precise neuronal states, and there are reports of modifications to the Ngn2-iN protocol where adding developmental patterning factors to the culture media can steer neuron differentiation to a desired path (Chen et al., 2020; Nehme et al., 2018). Our data supports a continued effort into identifying combinatorial transcription factor overexpression systems (Ravasi et al., 2010) and media conditions that can support precise neuron cell type engineering. Furthermore, comprehensive human nervous system reference cell atlases are required to understand the identity of cell states that emerge in *in vitro* engineered neuron systems. Single-cell genomics and comparisons to high-dimensional reference atlases should become a gold-standard to assess the heterogeneity and precision of *in vitro* engineered cells.

## ACKNOWLEDGEMENTS

We thank the Camp and Treutlein labs for helpful discussions. JGC and BT are supported by the Chan Zuckerberg Initiative DAF (grant number CZF2019-002440), an advised fund of Silicon Valley Community Foundation. JGC is supported by the European Research Council (Anthropoid-803441) and the Swiss National Science Foundation (Project Grant-310030_84795). BT is supported by the European Research Council (Organomics-758877, Braintime-874606), the Swiss National Science Foundation (Project Grant-310030_192604) and the National Center of Competence in Research Molecular Systems Engineering.

## AUTHOR CONTRIBUTIONS

MSc established the Ngn2 iPSC lines. MSc and SE generated iN cultures with assistance from AW. SE and WH established the selective single cell dissociation. SE generated the time course and d35 scRNA-seq Ngn2-iN scRNA-seq data with support from MSc, MSa, and WH. MSc with support from SE and AW generated IHC data. NNK provided guidance to establish the Ngn2 iPSC lines. ET provided guidance for IHC and iN culture. HCL generated the Ngn2 induction time course scRNA-seq data with support from SE. ZH, SE, and HCL analyzed the scRNA-seq data. HCL, ZH, SE, BT, and JGC designed the study and wrote the manuscript.

## DATA AVAILABILITY

The processed single-cell RNA-seq data and computaitnal codes used in this study were deposited on Mendeley Data with doi: 10.17632/y3s4hnyvg6.

## Experimental and computational procedures

### Cell culture

All cells described in this work were incubated at 37 °C, 5% CO2 and 90% humidity unless otherwise stated. 409b2 (RIKEN BRC cell bank), Sc102a1 (System Biosciences) stem cells and corresponding rtTA/Ngn2-derivatives were cultured in standard feeder-free conditions in mTeSR1 (StemCell Technologies) on plates coated with matrigel (Corning). Primary cortical rat astrocytes (Gibco) were cultured in high-glucose DMEM containing 10% FCS and 1% Pen/Strep on plates coated with poly-D-Lysine (Sigma-Aldrich). The astrocytes were fed every 4-5 days and passaged once a week with standard Trypsin-EDTA digestion.

### Generation of rtTA/Ngn2-iPSCs

rtTA/Ngn2 double positive stem cell lines were generated as previously described (Frega et al., 2017). Briefly, 409b2 or Sc102a1 stem cells were transduced with lentiviruses carrying rtTA (reverse tetracycline-controlled transactivator)-Neo cassettes and Ngn2-Puro cassettes. Transduced cells were selected for stable double integration using G418 and Puromycin treatment. rtTA/Ngn2-409b2 cells were either propagated as a non-clonal pool or passed selection by picking a colony derived from a limiting dilution of a single-cell suspension to form a clonal line (monoclonal 409b2). The resulting 409b2, Sc102a1 and monoclonal-409b2 rtTA/Ngn2-derivatives were subsequently used for differentiating Ngn2-iNeurons.

### Differentiation of Ngn2-iNs from iPSCs

rtTA-Ngn2-iPSCs were differentiated into Ngn2-iNs as previously described (Frega et al., 2017). To differentiate rtTA-Ngn2-iPSCs into Ngn2-iNs, rtTA-Ngn2-iPSCs were seeded as single cells on plates coated with 50 μg/mL poly-L-ornithine in borate-buffer and 10 μg/mL Laminin in DMEM/F-12, with Ngn2 expression induced using 4 μg/mL Doxycycline. One day after seeding, the medium was changed to DMEM/F-12 with 4 μg/ml Doxycycline, 1:100 N-2 supplement, 1:100 MEM non-essential amino acid solution (NEAA), 10 ng/μL human BDNF, 10 ng/μL human NT3, 1:1000 ROCK inhibitor and laminin 0.2 μg/mL. Two days after seeding, rat astrocytes were added to the culture. On the third day after seeding, the medium was changed to Neurobasal medium with 4 μg/ml Doxycycline, 1:50 B-27 supplement, 1:100 glutamax supplement, 10 ng/μl human BDNF, 10 ng/μl human NT3 and 2 μM Cytosine-D-arabinofuranoside (Ara-C). On day 5, 7 and 9 after seeding, 50% of the medium was exchanged with Neurobasal-based medium similar as day 3 only without the Ara-C. After day 9, 50% of the medium was changed every second day using the Neurobasal-based medium containing 2.5% FCS. Ngn2-iNs were kept for the indicated time and dissociated for scRNA-seq.

### Immunostaining of iNeurons

iNeurons, grown on coated coverslips (acid-treated, Kleinfeld Labortechnik), were fixed with 2% PFA (prepared freshly, pre-warmed to 37 °C) for 8 min. The coverslips were washed three times with 2 mL PBS and stored in PBS at 4 °C. iNs were permeabilized with 0.05% Triton X-100 in PBS for 10 min and quenched in 1 ml 0.2 M Glycine buffer for 30 min at room temperature. The coverslips were washed with PBS and incubated with 100 μl of the respective primary antibody diluted in IF-buffer (20 mM phosphate buffer cointaining 0.2% gelatin and 0.05% Triton X-100). The coverslips were washed with IF-buffer five times for 5 min and then incubated with 100 μl of the respective secondary antibody diluted in IF-buffer containing DAPI (dilution 1:1000). After five washes with IF-buffer for 5 min and three quick washed with PBS, the coverslips were mounted on slides with Mowiol 4-88 and stored at 4 °C.

iNs were acquired as confocal Z-stacks. iNs were imaged using an Olympus FV1200 confocal microscope equipped with a 10-x objective or 60-x oil immersion objective (optical section thickness: 1.028 μm, distance between consecutive optical sections: 0.4 μm). We acquired three-dimensional (Z-stack) scans with a number of z-sections ranging from 5 to 30 depending on the cell. Single tiles were 1024 × 1024 pixels. The following lasers were used: 405 nm for DAPI, 488 nm for MAP2/TUJ1.

### Dissociation of Ngn2-iNs for scRNA-seq

To prepare single-cell suspension for scRNA-seq, Ngn2-iNs were selectively dissociated from co-cultured rat astrocytes using a mild dissociation procedure. In this procedure, cultured Ngn2-iNs were washed with DPBS and incubated at 37 °C with a 1% Accutase in EDTA solution for 5 minutes. The digestion was then neutralized using Neurobasal medium with 10% FCS. The media was gently flushed onto the cells using a wide bore tip. The progress of selective neuron detachment was constantly monitored under the microscope. Enriched neural cell suspensions were collected in 15 ml tube and centrifuged for 5 minutes at 300 g. The cell pellets were resuspended and further dissociated in undiluted Accutase under 5-10 minutes incubation at 37 °C. The dissociated cells were neutralized, centrifuged, resuspended in the residual 100 μl medium, triturated 20 times, strained using a 30 μm pre-separation filter and quantified using a Countess automated cell counter. Cell suspensions were then diluted to 500-1000 cells/mL for subsequent scRNA-seq experiment.

### scRNA-seq library preparation and sequencing

For scRNA-seq library preparation from aforementioned single-cell suspensions, Chromium™ Single-cell 3’ Reagent Kits (10xGenomics, Pleasanton, USA) were applied according to manufacturer instructions. The Chromium™ Single-cell 3’ Reagent Kits v2 was employed on Ngn2-iN generated from 409b2 time-course experiments, monoclonal 409b2 and Sc102a1 iPSCs with approximately 3,000 cells loaded per lane on a 10x microfluidic chip device. Chromium™ Single-cell 3’ Reagent Kits v3 was used on Dox-treatment duration experiments with nearly 8,000 cells loaded per lane. Quantification and quality control of the 10x library was carried out on Bioanalyzer (Agilent) using High Sensitivity DNA chips. Libraries prepared from the 10x v2 kit and v3 kit were respectively sequenced on the Illumina HiSeq 2500 and Illumina NovaSeq S1 platform.

### Preprocessing of raw sequencing data

We used Cell Ranger, the suggested analytic pipeline by 10x Genomics, to demultiplex raw base call files to FASTQ files and align reads to the mouse-human dual species genome and transcriptome (mm10-hg38, provided by 10x Genomics) with the default alignment parameters. Demultiplexing of human and mouse cells was done based on their read mappability to the two genomes by Cell Ranger. Only human cells were kept for the following analysis. Cell Ranger was used again to map the human reads to the human-only genome and transcriptome (hg38, provided by 10x Genomics). Pooled samples of different human lines (409b2 and Sc102a1) were demultiplexed using demuxlet (Kang et al., 2018), based on the genotyping information of the two lines.

To quantify expression levels of the exogenous Ngn2 in cells, the unmapped reads after mapping to the human genome were extracted and compared to the sequences of the inserted Ngn2. In brief, for each read a series of substrings were obtained with a sliding window size of 40nt and step size of 10nt. The read was considered as a hit of exogenous Ngn2 transcript if at least 50% of the read substrings were perfectly matched to the inserted Ngn2 transcript sequence.

### Analysis of the time course scRNA-seq data of Ngn2-iN induction

Seurat (v3.1) was applied to the scRNA-seq data for further preprocessing. Quality control (QC) was done by excluding cells with more than 6000 genes detected, as well as those with mitochondrial transcript proportion larger than 10%. A subset of cells from the iPSC samples and those at day 1, 5, 14 and 29 since doxycycline treatment, which have detected gene number between 1500 and 5500 as well as mitochondrial transcript proportion less than 7% was further extracted. Highly variable genes were identified using the ‘vst’ method, and louvain clustering (resolution = 1) was used to do clustering to the cells. A k-nearest neighbor (kNN) graph was generated and visualized by using SPRING (Weinreb et al., 2018).

To observe the molecular changes along the whole time course of Ngn2-iN induction process from iPSC more comprehensively, we applied Cluster Similarity Spectrum (CSS) (He et al., 2020) to all the cells passing the initial QC, to reduce batch effect between samples of different time points. In brief, 5,000 highly variable genes were firstly identified using the ‘vst’ method. Their expression levels across cells were z-transformed, followed by principal component analysis (PCA) for dimension reduction. Next, cells from each sample were subset, and louvain clustering (resolution = 0.6) was applied based on the pre-calculated top-20 PCs. Average expression of the pre-defined highly variable genes was calculated for each cluster in each sample. Spearman correlation coefficient was calculated between every cell and every cluster in different samples. For each cell, its correlations with different clusters of each sample were z-transformed. Its z-transformed similarities to clusters of different samples were then concatenated as the final CSS representation. Louvain clustering (resolution = 0.5) was applied to the CSS representation.

Next, pseudocells were constructed, using a similar method as described (Kanton et al., 2019). In brief, cells from the same sample and assigned to the same cluster were grouped into kNN territories, each of which was centered by one randomly selected cell with its kNNs (k=20) surrounded. The CSS representation of each pseudocell was calculated as the average CSS representation of its included cells. A constrained kNN network (k=20) of pseudocells was constructed, to only consider pseudocells from the same or nearby time points when screening for nearest neighbours. The kNN network was then visualized by SPRING (Weinreb et al., 2018). The same projection method based on the support vector regression model as described (Kanton et al., 2019) was applied to project the single-cell data to the cell embedding space that was defined for pseudocells.

To better illustrate temporal molecular changes along the Ngn2-iN induction process, pseudotime analysis was applied to the CSS-represented time course scRNA-seq data using the diffusion map algorithm (implemented in the R package destiny, k=50). Cells in two side-branch clusters (Cl-8, Cl-15) were excluded from the pseudotime analysis. The ranks of DC1 were used as the pseudotimes. An F-test-based test was applied to the expression profile along pseudotimes to identify genes with pseudotime-dependent expression changes. In brief, for each gene, a natural spline linear regression model (df = 5) was constructed for cells along the pseudotime course. The residuals of the variation, which cannot be explained by the model, were then compared to the total variation of the gene by an F-test (Bonferroni corrected P<0.01). A kNN network (k=50) was constructed for the identified genes, based on the pairwise Pearson correlation distances. Jaccard indices between genes were calculated based on the resulting kNN network to weight the network, and edges with Jaccard indices less than 1/15 were pruned. UMAP and louvain clustering (resolution = 0.5) was applied to the weighted gene kNN network to construct the gene embeddings and identify gene clusters.

To infer the gene regulatory network (GRN) that contributed to the Ngn2-iN induction initialization, cells at the samples at induction time point h6-12 and d1 were subset. The pySCENIC pipeline (Aibar et al., 2017) was applied with default settings except for allowing inhibitory regulons to be identified. Transcription factor network was derived from the resulting regulons and the igraph package in R was used to generate the layout of the network for visualization.

### Analysis of the scRNA-seq data of Ngn2-iN of multiple cell lines at week5

The scRNA-seq data of Ngn2-iN cells at day 35 of samples from the 409b2 and Sc102a1 human iPSC lines, including those from the monoclonal 409b2 iPSC cells, were log-normalized using Seurat (v3.1). 3,000 highly variable genes were identified for cells from 409b2, 409b2 monoclonal and Sc102a1 iPSCs separately. The data was integrated with Seurat. UMAP and louvain clustering (resolution = 0.3) was applied to the integrated data. During this process, the first 20 components of canonical component analysis (CCA) and PCA were considered. Two resulting clusters (Cl-0 and Cl-1) were merged as no strong signature unique to one of the two clusters was observed. Cluster markers were identified using the R package presto, defined as genes that being detected in at least 50% of cells in the cluster and at least 20% higher than in other cells, with AUC>0.7, expression fold change > 1.2 and Benjamini-Hochberg (BH) corrected two-sided Wilcoxon’s rank sum test P<0.01.

To characterize each Ngn2-iN cluster, we retrieved the scRNA-seq data of three mouse reference data sets: the adolescent mouse nervous system atlas (Zeisel et al., 2018) which includes both the central nervous system (CNS) and peripheral nervous system (PNS) in adolescent mice; the developing mouse brain (La Manno et al., 2020); and the developing mouse retina (Clark et al., 2019). Top-5000 highly variable genes in the adolescent mouse nervous system and the developing mouse brain were identified using the vst method in Seurat and intersected, resulting in 1952 shared variable genes with one-to-one human orthologs according to Ensembl (v84) annotation. Average expression profiles were calculated for each of the annotated neuron subtypes in the three atlases, as well as the 5 Ngn2-iN clusters, across the shared variable genes. Pearson correlation coefficient was then calculated between average expression profiles of Ngn2-iN clusters and selected mouse reference neuron subtypes, including 14 subtypes in developing brain, the developing spinal cord motor neuron, 6 subtypes in developing retina, and 6 subtypes in adolescent PNS.

To further benchmark the Ngn2-iN with other iPSC-derived neurons, we additionally retrieved the scRNA-seq data of human cerebral organoids (Kanton et al., 2019) and retinal organoids (Cowan et al., 2019). Similarly, average expression profiles of the seven neuron subtypes in the cerebral organoids, as well as those of the five neuron subtypes in the retinal organoids, were calculated across the shared variable genes obtained above. The resulting profiles were correlated with the mouse references similar to the Ngn2-iN clusters.

To summarize the CNS/PNS signatures in the Ngn2-iN cells, a transcriptome deconvolution based on quadratic programming (Treutlein et al., 2016) was used to dissect the CNS and PNS neuronal signatures. In brief, differential expression analysis using the presto package was applied to compare transcriptome of CNS and PNS neurons in the mouse nervous system, based on the adolescent mouse brain atlas mentioned above. Differentially expressed genes (DEGs) were identified as genes with BH corrected P<0.01 and expression fold change > 1.2. The average expression levels of those CNS-PNS DEGs were calculated for each annotated cell type in the mouse nervous system (annotation layer 4). Considering the transcriptome of each Ngn2-iN cell as a mixture of transcriptomes of different cell types, quadratic programing was then used to calculate the relative contribution of each cell type by solving the constrained linear least-square problem:

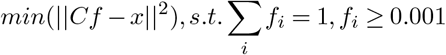

Here *f* is the vector of relative cell type contribution, *C* is the gene expression matrix of the CNS-PNS DEGs in annotated mouse cell types, and *x* is the expression level of these genes in a cell. Afterwards, the contributions of all CNS neuron subtypes were summed up as the CNS signature score of a cell.

To compare Ngn2-iN cells with primary fetal and adult neurons, the Fluidigm C1-based single cell RNA-seq data of human fetal and adult cortex was retrieved from SRA (SRP057196) (Darmanis et al., 2015). Average expression levels of each annotated major cell type, including replicating cells (neural progenitor cells) and quiescent cells (immature neurons) in fetal samples, as well as cell types in adult samples (neurons, astrocytes, oligodendrocytes, oligodendrocyte precursor cells, microglia, endothelial cells). Pearson correlations were calculated across expression of highly variable genes in the Ngn2-iN d35 data set, between each Ngn2-iN cell and each annotated cortical major cell type.

### Analysis of the scRNA-seq data of varied doxycycline treatment duration

The Ngn2-iN scRNA-seq data with varied doxycycline treatment durations was preprocessed using Seurat similar as above. As quality control, only cells with detected gene numbers between 1370 and 8000 remained in the following analysis. The lower bound of the detected gene number was the minimum threshold to exclude cells with detected gene numbers being at the lower peak of the observed bimodal distribution of detected gene numbers in cells. CSS representation was then calculated to integrate cells from different Ngn2-iN samples with varied doxycycline treatment duration (cluster resolution = 1). UMAP and louvain clustering (resolution = 0.3) was applied to the CSS representation of cells.

## Supplementary Figures

**Supplementary Fig. 1.**
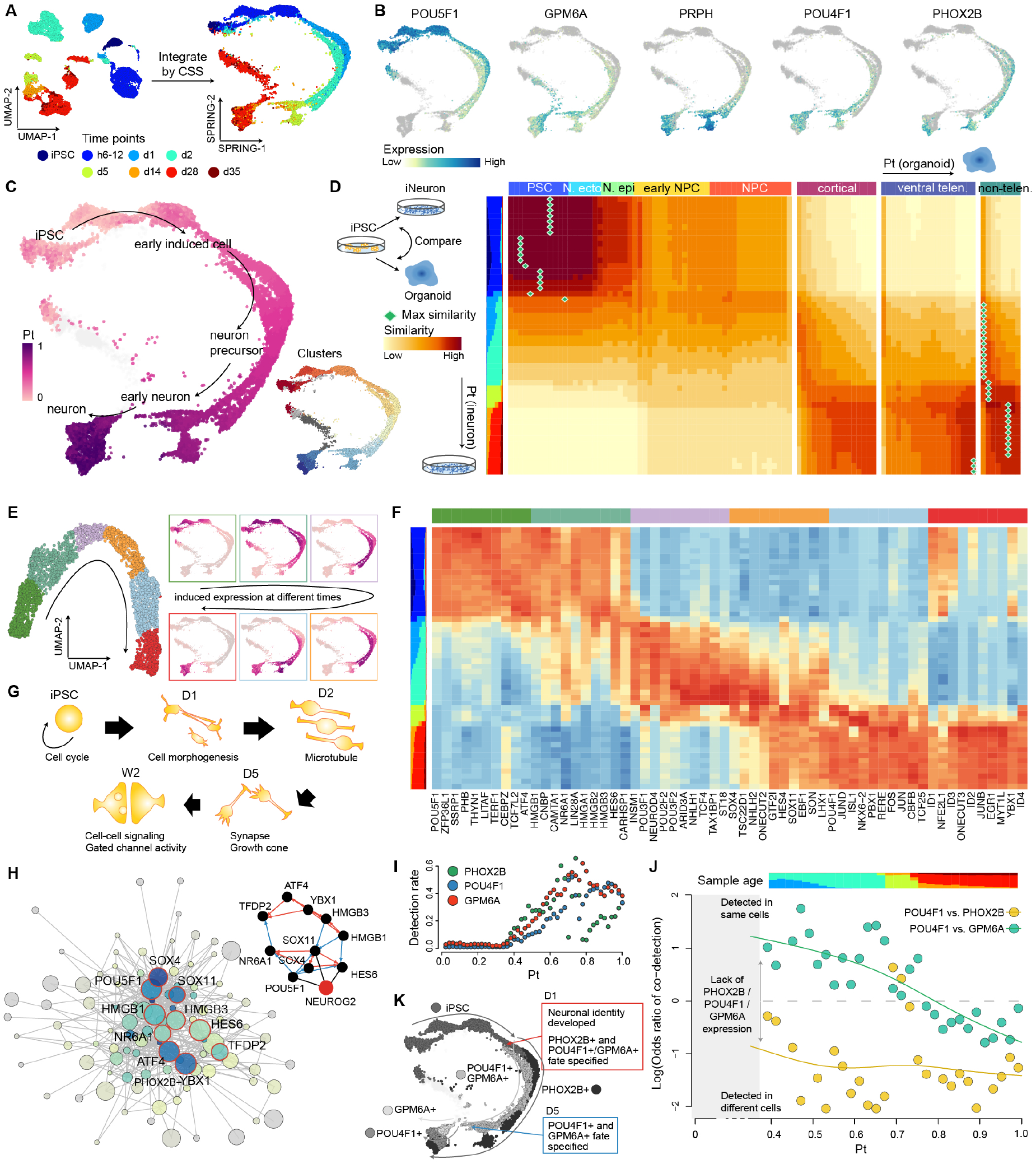
Analysis of Ngn2-mediated direct reprogramming path from iPSC to induced neurons. (A) Ngn2-iNs directly reprogrammed from 409b2 iPSCs were harvested at different time points after induced expression of Ngn2 by Dox treatment and analyzed by scRNA-seq. scRNA-seq data was directly analyzed by UMAP or combined by CSS-based integration with differentiation trajectory reconstructed using SPRING. UMAP and SPRING plots were colored by time points. h6-12 is the 1:1 mixture of cells harvested after 6h and 12h Dox induction, respectively. (B) SPRING plots colored by marker gene expression. (C) Each cell in the Ngn2-iN reprogramming path was ordered in pseudotime based on transcriptome similarities. The derived pseudotime was then colored on the SPRING plot. Cell clusters along the developmental trajectories were also color labeled. (D) Developmental trajectories from iPSCs to Ngn2-iNs and brain organoids (Kanton et al., 2019) were aligned and compared. (E) 3,231 genes among the 17,198 genes detected in more than 50 cells along the full time course of Ngn2-iN development had significant change in expression. These genes were grouped into 6 clusters with their expression colored on the SPRING plot. (F) Heatmap of transcription factors (TFs) in each cluster among the differentially expressed genes identified in (E). (G) Schematic representation of the biological processes identified in the functional enrichment analysis from differentially expressed genes in (E). (H) Gene regulatory networks for TFs with pseudotime-dependent expression. Highlighted are the top-10 TFs with highest expression and most connection with other TFs. (I) Detection of cells with PHOX2B, POU4F1 or GPM6A expression along the pseudotime of Ngn2-iN development. (J) Coexpression of POU4F1/PHOX2B and POU4F1/GPM6A along the pseudotime of Ngn2-iN development. (K) Fate specification of the three Ngn2-iN neural subtypes along the time course of development. The SPRING plot was colored by cell identity.

**Supplementary Fig. 2.**
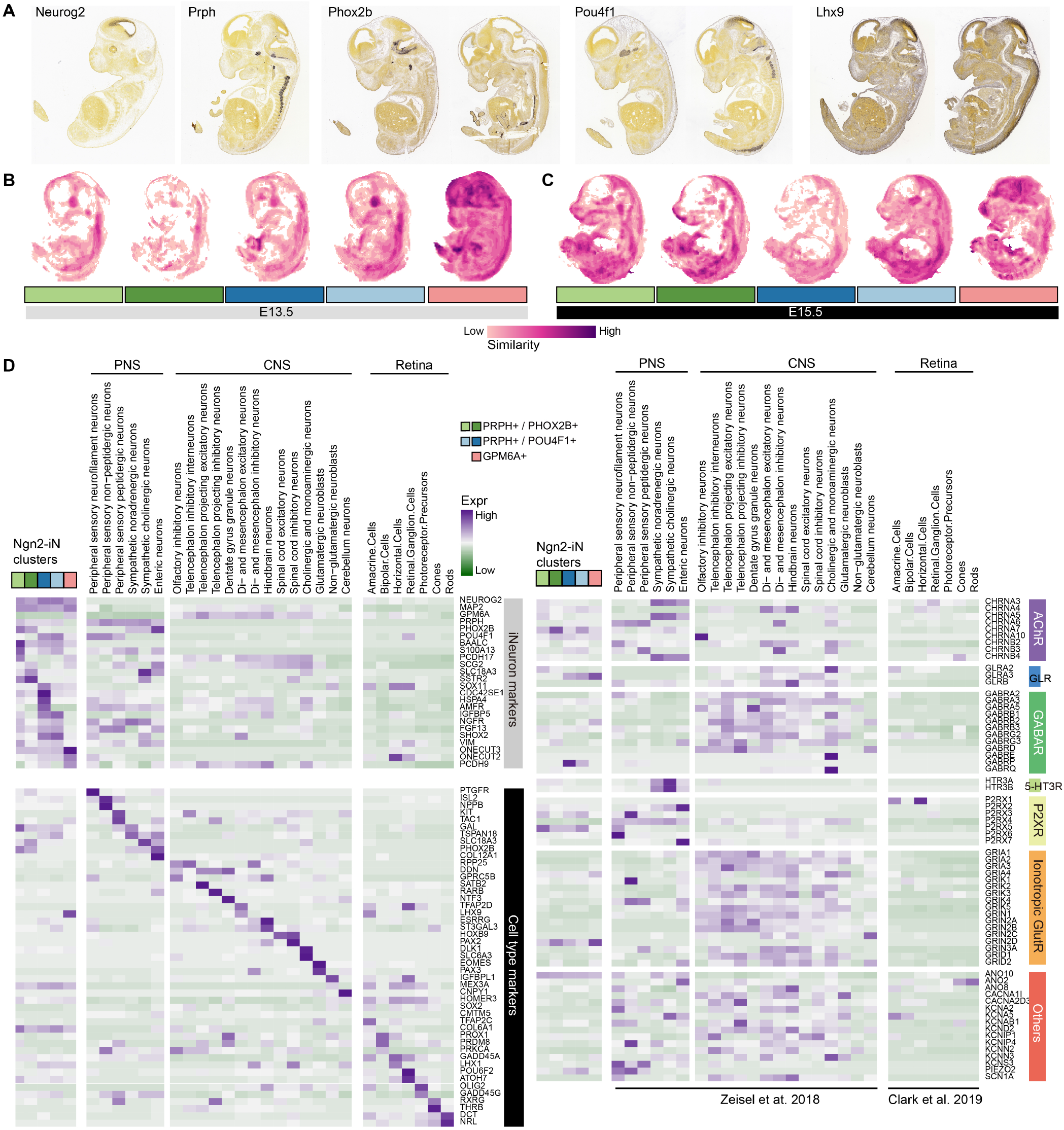
Transcriptomic comparison of Ngn2-iN clusters to primary mouse references. (A) Spatial expression patterns of selected markers via in situ hybridization (ISH) in the E13.5 mouse embryos from the Allen Developing Mouse Brain Atlas. (B-C) Transcriptomic similarity between each of the five Ngn2-iN populations and E13.5 (B) and E15.5 (C) mouse embryos. It shows the maximum similarity projections across sagittal sections in the mouse embryos from the Allen Developing Mouse Brain Atlas. (D) Average expression of various marker genes Ngn2-iN clusters and primary mouse PNS and CNS neuron subtypes (left), as well as various genes encoding ion channels (right), in Ngn2-iN clusters as well as primary mouse neuron subtypes.

## Supplementary Tables

**Table S1.** GO terms enriched in the course of Ngn2-iN development

